# Genomic-island cassette architecture drives pathogenic *Enterococcus cecorum* lineages: Cassette2Vec-EC, a structural genomics and machine-learning framework

**DOI:** 10.64898/2026.02.21.707173

**Authors:** Rushikesh R. Lagad, Shakil Rafi, Aranyak Goswami

## Abstract

Mobile genetic elements and genomic islands (GIs) frequently encode antibiotic resistance and host-adaptation cargo, yet routine genome comparison pipelines often miss the higher-order organization of how genes co-occur as transferable, GI-anchored modules. We present Cassette2Vec-EC, a structural genomics framework that converts annotated genomes into cassette units (local gene neighborhoods with GI context), encodes each cassette as a fixed-length feature vector, and applies genome-grouped machine learning to predict pathogenic lineages while preventing within-genome leakage. Using a curated *Enterococcus cecorum* cohort from poultry production systems, we integrate pangenome context, GI calls, mobility markers, and AMR/virulence annotations into cassette-level features and evaluate models strictly under GroupKFold-by-genome. Cassette2Vec-EC achieves strong genome-level generalization (AUROC 0.975 ± 0.030, average precision 0.938 ± 0.077, Brier score 0.056 ± 0.058). When evaluated at the cassette unit level under the same genome-grouped protocol, performance remains high (AUROC 0.974 ± 0.029, AP 0.919 ± 0.093, Brier 0.057 ± 0.057), supporting that cassette representations capture transferable signal rather than genome identity. Baselines show that GI burden alone can partially rank genomes but yields poorer calibration and limited interpretability. By combining comparative genomics with cassette-aware features and providing locus-level explanations (SHAP) that map predictions to specific GI-associated modules, Cassette2Vec-EC provides a practical blueprint for genomic-island–aware pathogen surveillance, junction-based diagnostics, and targeted monitoring of high-risk lineages.

**Importance:** Pathogenic *Enterococcus cecorum* lineages threaten poultry health and production, yet many genomic surveillance workflows reduce genomes to gene presence–absence lists and overlook how mobile elements are organized into transferable modules. Cassette2Vec-EC addresses this gap by representing genomic islands as GI-anchored cassette neighborhoods and encoding each cassette as a fixed-length feature vector linking cassette architecture to pathogenic risk. Under a strict genome-grouped evaluation protocol, the framework predicts pathogenic lineages while preventing within-genome leakage and supports calibrated probability estimates in the full model. Cassette level explanations (SHAP) localize predictive signal to specific GI-anchored modules, yielding tractable targets for surveillance prioritization and junction-based diagnostics. Although developed for poultry-associated *E. cecorum*, this cassette and GI-aware representation is transferable to other bacterial pathogens to strengthen genomic surveillance and help partners identify high-risk isolates before clinical disease emerges.

## Introduction

*Enterococcus cecorum* (EC) has emerged as an important poultry-associated pathogen (1, 2), with outbreaks linked to osteomyelitis, lameness, and production losses (3–6). Molecular epidemiology and diversity studies have documented multiple circulating EC lineages and outbreak-associated clones, underscoring the value of molecular typing for surveillance (7–9). Phenotypic characterization and environmental persistence work further support EC’s capacity to persist and disseminate in poultry environments (10, 11). Comparative genomics has revealed extensive accessory genome variation in EC, including intercontinental spread of pathogenic lineages (12–14). Many clinically relevant traits are encoded on mobile genetic elements particularly genomic islands (GIs) that package resistance, virulence, and mobility functions into transferable modules (13, 15–18). Standard gene-presence workflows treat these loci as independent features, often losing information about neighborhood structure, genomic context, and transferability.

Systems microbiology increasingly emphasizes higher-order genomic organization modules, neighborhoods, and transferable units rather than isolated loci. In this study, we operationalize that idea by defining cassette units as local gene neighborhoods with GI context and encoding each cassette into a fixed-length feature vector summarizing island geometry, mobility markers, and cargo signals. We then evaluate prediction of pathogenic lineages under a genome-grouped protocol (GroupKFold-by-genome) to ensure that no cassette rows from the same genome appear in both training and test sets (19). This paper focuses on three questions: (i) is pathogenicity signal explained by simple GI burden, or by specific cargo modules; (ii) can a cassette- and GI- aware representation improve predictive performance and calibration; and (iii) can model explanations be mapped back to biologically meaningful, transferable cassette modules that enable translational surveillance and diagnostics.

Beyond gene presence, EC adaptation is shaped by how accessory genes are organized into transferable units. Horizontal gene transfer in *Enterococcus* frequently moves clusters of resistance, virulence, and mobility functions as physically linked blocks on genomic islands and other mobile elements (13, 18). This modular architecture motivates a structural-genomics view: pathogenic potential may depend less on any single locus and more on recurrent, island-anchored cassette modules that co-occur and move together.

Accordingly, we treat cassette neighborhoods as the unit of learning and use cassette-level performance as a mechanistic check that the representation captures modular signal. The primary objective, however, remains genome-level risk prediction under a strict genome-grouped protocol that prevents within-genome leakage and supports surveillance-oriented generalization.

## Materials and Methods

### Genome cohort, phenotype labels, and annotation harmonization

We analyzed a curated *Enterococcus cecorum* (EC) genome cohort derived from poultry production systems, with isolates labeled as commensal or pathogenic based on sampling context and phenotype metadata (Supplementary Data S1). Genomes were uniformly annotated using Prokka (20). Genomic islands (GIs) were predicted using IslandViewer 4 (21). Antimicrobial resistance (AMR) loci were screened using ABRicate (22) against curated resistance databases including ResFinder (23) and CARD (24). Orthologous clustering was performed with PIRATE (25) to stabilize gene block assignments across assemblies and support neighborhood-based cassette construction. Functional annotation of non-AMR GI genes was performed using eggNOG-mapper v2 (26) to provide COG functional categories and KEGG pathway assignments for biological interpretation. eggNOG-mapper outputs were used for mobility marker keyword parsing and non-AMR mobilome signature annotation; the corresponding schema fields (eggNOG_class, pathway) are included in the v1 feature matrix as reserved columns but were not populated or used as direct model inputs in the analyses reported here (see feature schema note above).

Operational label definition. We operationally defined “pathogenic” isolates as those recovered from clinical disease contexts consistent with EC-associated pathology in poultry production systems (e.g., lesion-associated clinical submissions such as osteomyelitis/lameness contexts and outbreak-linked cases when explicitly indicated), and “commensal” isolates as those derived from healthy-bird surveillance or non-disease-associated sampling contexts (e.g., routine monitoring without reported clinical disease). Because these labels are metadata-derived and EC pathogenicity can be context-dependent, we interpret phenotype labels as surveillance-grade and prioritize prospective validation to quantify performance under real-world sampling variability.

### Genomic island calling and interval harmonization

Genomic islands were predicted for all genomes using IslandViewer 4 (21), which integrates sequence composition and comparative genomics signals to identify horizontally acquired regions. To focus on structurally meaningful mobile regions rather than isolated genes, we retained predicted islands ≥5 kb.

To address overlapping and nested IslandViewer calls within a genome, we collapsed predicted GI intervals to a non-redundant merged set prior to cassette construction. Two intervals were considered overlapping if their closed coordinate intervals shared at least one basepair of overlap (intersection length ≥1 bp; i.e., max(start1,start2) ≤ min(end1,end2)), such that “touching” at a single base was treated as overlap. Overlapping/nested intervals were merged to their union using standard interval-union logic: intervals were sorted by start coordinate and iteratively merged by extending the current interval’s end coordinate while overlap held; otherwise, the current merged interval was emitted and a new interval was started. A gene was labeled GI-associated if any portion of its annotated span overlapped a merged GI interval (≥1 bp overlap).

### AMR and mobilome feature derivation

AMR loci were identified using ABRicate (22) against ResFinder (23) and CARD (24). Genes matching AMR hits were flagged as AMR-positive for feature construction and were excluded from the non-AMR mobilome signature derivation to avoid conflating resistance cargo with non-AMR mobilome features.

Mobility-associated genes were identified using standardized annotation flags and/or keyword/domain-informed parsing applied to Prokka and eggNOG outputs (20, 26). Mobility markers included integrase, transposase/IS, and recombinase annotations. These mobility markers were incorporated as individual indicators and as an aggregated mobility-burden feature at the cassette scale (see below).

### Cassette construction and GI-anchored neighborhood definition

Cassette units were defined as contiguous gene neighborhoods (operon-scale runs of PIRATE ortholog blocks) located within predicted genomic islands (21), bounded by PIRATE-defined ortholog blocks (25) and merged genomic-island intervals. In the primary analysis, cassettes were GI-anchored and therefore did not intentionally span non-GI regions; genes outside predicted islands were used only to construct non-GI neighborhoods in baseline/ablation analyses.

Cassette boundary definition (operational). For each genome, Prokka annotated genes were ordered by contig and genomic coordinate. Within each merged GI interval, we extracted the ordered list of GI-associated genes and defined a cassette neighborhood as a maximal contiguous run of GI-associated genes along the coordinate axis. Contiguity was broken when (i) the next gene fell outside the merged GI interval, (ii) the contig ended, or (iii) a GI-associated run was interrupted by a non-GI gap. Large islands were not artificially capped by gene count; thus, a long GI remained a single cassette neighborhood unless interrupted by contig boundaries or non-GI gaps. These cassette neighborhoods constitute the fundamental tokens used for downstream modeling.

Cassette construction depends on IslandViewer predictions, the ≥5 kb island filter, and coordinate-based contiguity rules. While we use this primary operational definition for reproducibility, alternative definitions (e.g., different GI callers, different minimum GI length thresholds, or splitting long islands into multiple neighborhoods by gene-count or mobility-anchor boundaries) may alter cassette granularity and will be evaluated in sensitivity analyses.

### Cassette feature engineering: fixed-length numeric representation

Operationally, the Cassette2Vec representation was instantiated as a fixed-length numeric cassette feature vector encoding cassette structure and context (e.g., GI overlap/attributes and mobility/cargo indicators). This cassette-level vector served as the predictive model input, enabling genome-level risk prediction while preserving interpretability at the cassette/module level.

Full (all numeric) feature set. The Cassette2Vec-EC “Full (all numeric)” model uses 20 numeric input features (Table 2). To ensure reproducibility, we provide (i) a complete feature definition table listing all 20 feature names, types, units, allowable/observed ranges, and brief interpretations (Supplementary Data S4), and (ii) the released feature matrix schema used for modeling (cassette2vec_ML_features_v1_with_mobility_load.xlsx). Of the 20 declared schema fields, three (eggNOG_class, pathway, cluster_id) are reserved for forward compatibility and were not populated in the v1 feature matrix; all model performance metrics reported in Table 2 were obtained using the 17 active numeric inputs.

Feature schema note. The released v1 feature matrix includes three reserved schema fields (eggNOG_class, pathway, cluster_id) retained for forward compatibility; these fields are not populated in the v1 release and were not used as predictive inputs in the reported “Full (all numeric)” model. The complete set of model inputs is enumerated in Supplementary Data S4. Mobility-load score. Mobility-load was computed as a cassette-level mobility burden defined as the sum of mobility-associated gene indicators within the cassette neighborhood, based on integrase/transposase/recombinase classification (and additional mobility categories when applicable). Mobility-load therefore increases with the number of mobility-associated loci in a cassette and serves as a quantitative proxy for mobilome-rich, transfer-associated modules.

### Modeling framework and genome-grouped evaluation protocol

All predictive modeling was evaluated using 5-fold GroupKFold-by-genome (GroupKFold, n_splits = 5; scikit-learn [19]), ensuring that no cassette from a given genome appeared in both training and test partitions. This protocol prevents within-genome leakage and enforces generalization at the genome level. Because GroupKFold does not guarantee perfect class stratification, we report mean ± SD across folds, summarize fold-level class counts in the supplement, and interpret fold-to-fold variability in light of the limited pathogenic count per outer fold (∼10).

Genome-level performance was treated as primary: cassette-level predicted probabilities were aggregated per genome (mean predicted risk across cassettes) to yield a genome-level risk score used for pathogenic lineage classification.

### Model specification, training, and interpretability

We trained a gradient-boosted decision tree classifier implemented in XGBoost (27) on cassette feature vectors, producing cassette-level predicted probabilities that were aggregated to genome-level risk scores (mean predicted probability across cassettes per genome). All preprocessing steps were fit using only the training split within each outer fold and then applied to the held-out fold to prevent leakage.

Hyperparameter tuning was performed in a nested manner within the outer 5-fold genome-grouped evaluation: candidate settings were selected using genome-grouped splits on the training portion of each outer fold, then refit on the full outer training set prior to evaluation on held-out genomes. If class weighting was applied, it was computed within each training fold (e.g., scale_pos_weight) to avoid leakage.

For model interpretation, SHAP values were computed at the cassette-level (28). Genome-level explanations were obtained by aggregating cassette-level attributions across all cassettes within a genome, consistent with the probability aggregation used for genome-level risk scoring.

For exploratory visualization of cassette-vector structure, we trained a variational autoencoder (VAE) (29) and clustered embeddings using DBSCAN (30); these results are not shown but confirmed that cassette vectors form interpretable structure in latent space. VAE embeddings were used exclusively for visualization/clustering and were not incorporated as inputs to any downstream predictive model.

### Metrics, baselines, and signature stability

We report AUROC and average precision (AUPRC/AP) for ranking performance and Brier score (33) for probability calibration. To demonstrate that performance is not explained by trivial GI burden or AMR counting, we compared the full model against capacity-matched ablations/baselines (GI-only and mobilome-only) evaluated under the same genome-grouped protocol and tuned under comparable model-budget constraints.

Signature stability for the non-AMR mobilome panel was assessed by re-running the signature-selection procedure within each outer-fold training split and summarizing selection frequency across folds (Table S2; Supplementary Data S2).

### Non-AMR mobilome signature derivation and circularity control

The 20-gene non-AMR mobilome signature was derived using a training-only selection procedure to avoid circularity. Within each outer fold of the 5-fold genome-grouped evaluation, we computed gene-level enrichment statistics using only genomes in the training split. For each GI-localized non-AMR gene, enrichment was quantified as the log2 odds ratio of presence in pathogenic versus commensal genomes. Genes were ranked by absolute log2 odds ratio, and the top candidates were retained per fold.

The 20-gene non-AMR panel was derived within each outer-fold training split only; held-out folds were not used for selection.

Selection frequency across outer folds was computed to assess stability (Table S2). We report the panel as (i) a high-stability ‘core’ subset (selected in ≥3/5 outer training splits) and (ii) auxiliary loci retained to preserve family-level coverage and module completeness when multiple homologous/adjacent markers represent the same GI architecture. This avoids over-claiming per-locus stability while keeping the panel useful as a structural surveillance set.

This procedure ensures that the mobilome signature reflects reproducible pathogenic enrichment rather than artifacts of full-dataset overfitting.

## Results

### Cohort summary, cassette units, and GI coverage

We analyzed 145 *Enterococcus cecorum* genomes from poultry production systems (95 commensal and 50 pathogenic isolates; Supplementary Data S1). Across this cohort, cassette rows represent structured GI-anchored gene-neighborhood instances and serve as the unit of learning. IslandViewer outputs were available for the majority of genomes and were used for GI-burden and GI-context analyses.

### Assembly quality and potential confounding

Assembly fragmentation varied across the cohort (Supplementary Data S1). Commensal genomes had a mean contig count of 33.3 (median 33) and pathogenic genomes had a mean of 109.0 (median 72), computed across the full n = 145-genome cohort. Pathogenic assemblies showed substantially higher fragmentation on average, driven by 22 highly fragmented assemblies (>100 contigs; max 428 contigs), likely reflecting draft-quality sequencing from a subset of BioProject accessions; commensal assemblies showed no genomes exceeding 100 contigs. Because cassette continuity can be interrupted at contig boundaries, we performed a sensitivity analysis restricted to genomes with ≤50 contigs (high-contiguity subset). In the ≤50-contig subset (n = 91 genomes; 21 pathogenic; derived by filtering Supplementary Data S1 for assemblies with total_contigs ≤ 50), held-out genome-level performance remained high: AUROC = 0.982 (95% CI 0.955-1.000), AP = 0.935 (95% CI 0.858-1.000), and Brier = 0.031 (95% CI 0.006-0.061).

### GI burden is not the full pathogenicity signal

We first tested whether pathogenic genomes simply carry more predicted islands or greater total island length. Across available IslandViewer calls, GI count and total GI length show overlapping distributions between commensal and pathogenic classes (Figure 2; Table 1). Two-sided Wilcoxon rank-sum tests supported this overlap (GI count: p = 0.244, Cliff’s δ = -0.117; total GI length: p = 0.388, Cliff’s δ = 0.088), indicating that univariate GI-burden summaries do not separate classes.

**Figure 1.**
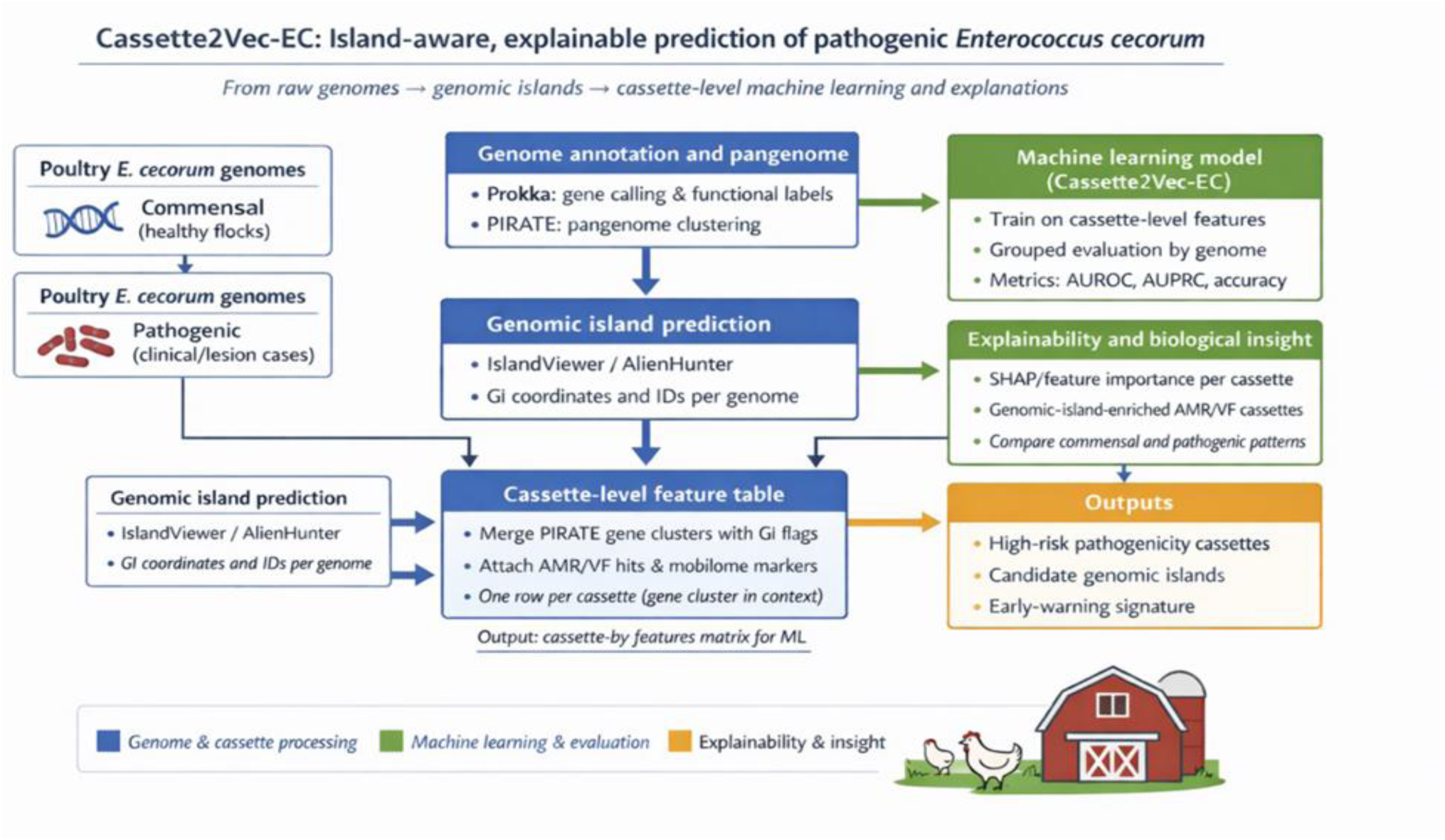
Cassette2Vec-EC workflow overview: genomes → GI prediction and annotation → cassette construction → cassette feature vectorization (Cassette2Vec encoding) → genome-grouped ML → SHAP-based interpretation.

**Figure 2.**
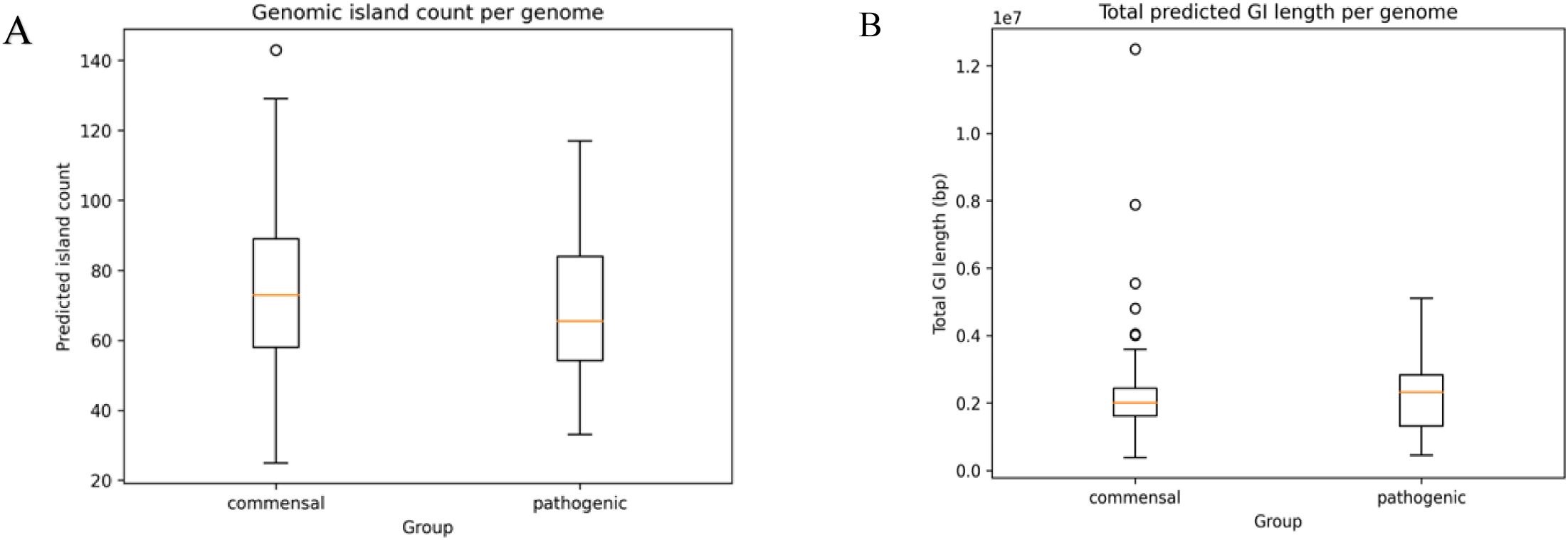
GI burden is not the full pathogenicity signal. **(A)** Predicted GI counts per genome. **(B)** Total predicted GI length per genome.

**Table 1.** Genomic island (GI) burden summary by class (IslandViewer-available subset). Values summarize predicted GI counts and lengths per genome; distributions are visualized in Figure 2.

| genome_class | n_genomes | median_islands | mean_islands | median_total_gi_length | mean_total_gi_length |
| --- | --- | --- | --- | --- | --- |
| commensal | 95 | 12 | 12.91 | 218622 | 225861 |
| pathogenic | 50 | 12 | 11.88 | 230631 | 238457 |

Per-genome genomic-island (GI) burden summaries (island counts and total island length per genome) are provided as Supplementary Data S3 (Supplementary_Data_S3_per_genome_GI_burden.xlsx).

Although total GI burden does not separate commensal and pathogenic genomes in univariate tests, GI-context features in aggregate still carry predictive information: the GI-only baseline achieves high ranking performance (AUROC = 0.963) but poor probability calibration (Brier = 0.215; Table 2), indicating that which islands and their contextual attributes matter more than island count alone.

**Table 2.** GroupKFold-by-genome performance for Cassette2Vec-EC and baseline feature ablations. Genome-level metrics are primary (mean ± SD across folds). GI-only ranking can be strong but calibration can degrade, motivating cassette/context modeling. †Three fields (eggNOG_class, pathway, cluster_id) are reserved schema placeholders and were not populated in the v1 feature matrix; the Full model used 17 active numeric inputs.

| Model | n_features | Genome AUROC (mean ± SD) | Genome AP (mean ± SD); Brier (mean ± SD) |
| --- | --- | --- | --- |
| Full (all numeric) | 20† | 0.975 ± 0.030 | 0.938 ± 0.077; 0.056 ± 0.058 |
| GI-only | 5 | 0.963 ± 0.047 | 0.911 ± 0.104; 0.215 ± 0.018 |
| Mobilome-only | 6 | 0.562 ± 0.055 | 0.432 ± 0.096; 0.232 ± 0.021 |

### Genome-grouped predictive performance and calibration

Under strict GroupKFold-by-genome evaluation, Cassette2Vec-EC achieves strong genome-level generalization (AUROC 0.975 ± 0.030; AP 0.938 ± 0.077; Brier 0.056 ± 0.058). Cassette-level metrics under the same genome-grouped protocol are similarly high (AUROC 0.974 ± 0.029; AP 0.919 ± 0.093; Brier 0.057 ± 0.057), supporting that predictive signal is not driven by within-genome redundancy.

The low Brier score of the full model indicates well-calibrated probability estimates across folds. In contrast, the GI-only baseline shows substantially poorer calibration, suggesting that island quantity alone provides ranking information but unreliable risk probabilities. Calibration curves (Figure 3C) show some fold-to-fold variability, likely reflecting limited per-fold pathogenic sample counts (∼10 per fold). For surveillance deployment, post-hoc recalibration (e.g., isotonic regression) and threshold selection would be advisable prior to operational implementation. Given that GroupKFold does not enforce class stratification, we report mean ± SD across folds and interpret variability in light of the limited pathogenic count per outer fold.

**Figure 3.**
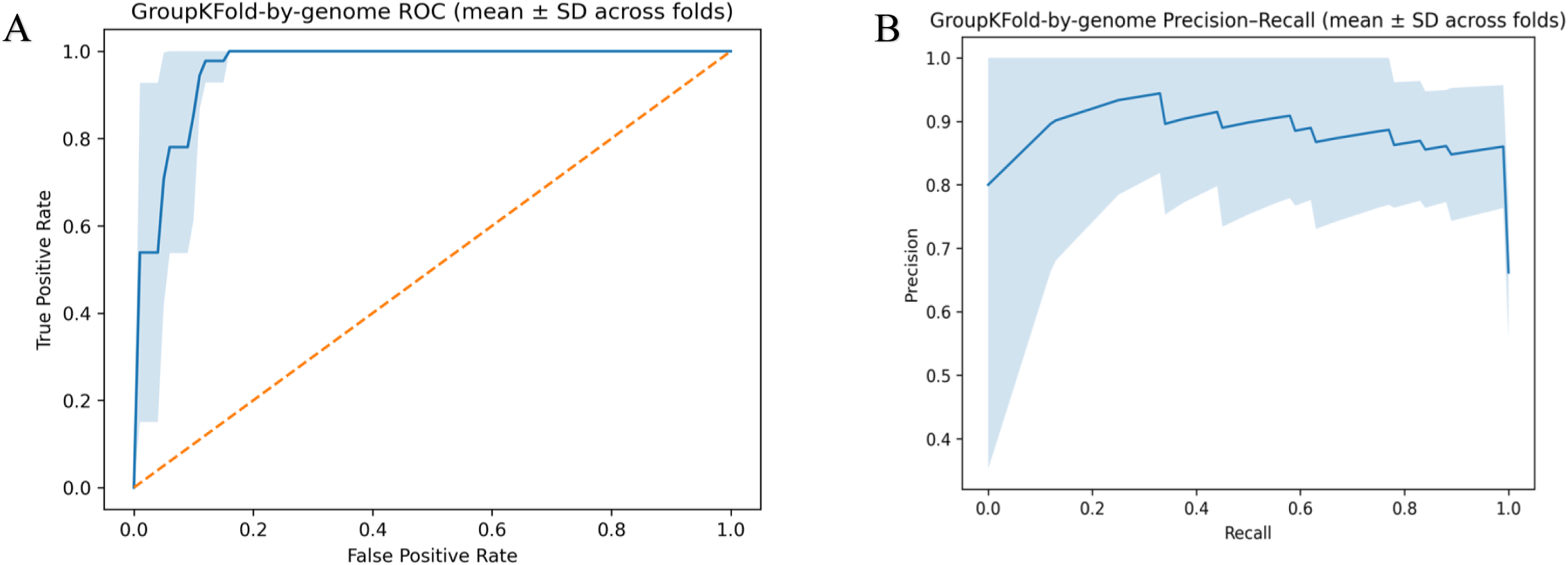
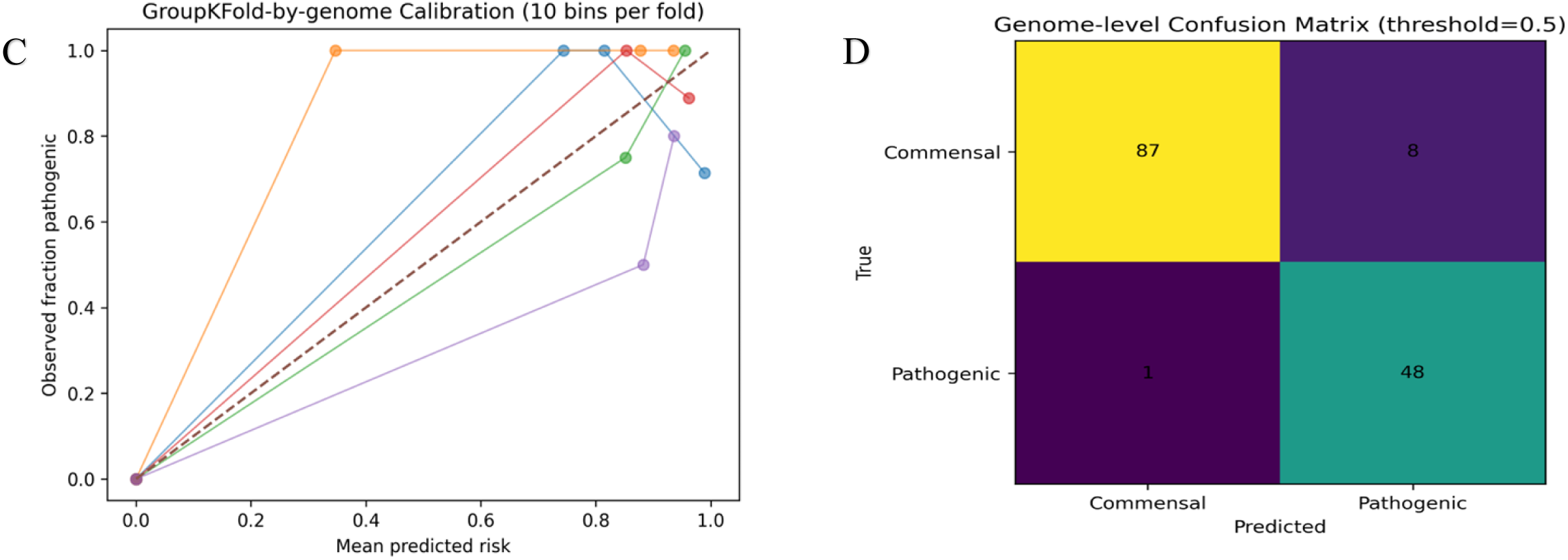
Genome-grouped predictive performance. **(A)** ROC curves across GroupKFold-by-genome folds. **(B)** Precision-recall curves across folds. **(C)** Binned calibration curves (10 bins per fold). **(D)** Genome-level confusion matrix at threshold 0.5.

### Baselines demonstrate signal beyond GI burden alone

Baseline comparisons (Table 2) show that GI-only features can rank genomes but provide limited interpretability and suboptimal calibration, while mobilome-only baselines underperform substantially. Together, these findings indicate that pathogenicity is not captured by island quantity or simple mobility counts alone; rather, multivariate cassette architecture—co-localized gene modules within GI context drives predictive performance and calibration.

### Derivation of a pathogenic-enriched non-AMR mobilome signature

To provide an interpretable structural panel beyond global GI burden, we derived a 20-gene non-AMR mobilome signature from recurrent GI-localized cassette modules enriched in pathogenic genomes. AMR loci identified by ABRicate were excluded from this panel by design to avoid resistance-driven confounding. AMR determinants were analyzed separately in the comparative AMR prevalence summary (Supplementary Figure S1). Notably, several non-AMR loci contribute to prediction, consistent with Cassette2Vec capturing GI architecture and ecological adaptation signals beyond antibiotic resistance.

Presence-absence patterns across genomes show marked class differences. Stability analysis across outer-fold training splits demonstrates consistent selection of several loci (Supplementary Table S2), supporting the robustness of the signature as a structural surveillance panel rather than a resistance-only marker set. The signature includes mobile-element recombination factors, plasmid replication and maintenance proteins, nucleotide and carbohydrate metabolism genes, and stress-response loci, providing biologically plausible candidates for host adaptation and persistence.

### Explainability links predictions to GI-anchored modules

SHAP analysis was performed at the cassette-row level to identify which cassette and context features drive pathogenic predictions. Cassette-level attributions were aggregated within genomes to support genome-level interpretation (Figure 4). The highest-impact features driving pathogenic predictions were amr_hit, Mobility_Load, and GI_AMR_density, each showing positive SHAP values concentrated in high-feature-value cassettes (Figure 4). Features reflecting mobility burden (transposase_flag, integrase_flag, recombinase_flag) contributed smaller but consistent positive signal, while AMR_neighborhood_score showed bidirectional effects, suggesting that AMR cargo spatial context modulates rather than uniformly increases predicted risk. These explanations enable actionable follow-up, including prioritizing surveillance of genomes enriched for recurrent high-risk cassette archetypes and designing junction-PCR assays targeting conserved cassette boundaries between mobility anchors and cargo loci. High-impact cassettes frequently combine mobility markers with putative host-interaction and metabolic genes, consistent with the hypothesis that recurrent GI-anchored modules contribute to pathogenic potential.

**Figure 4.**
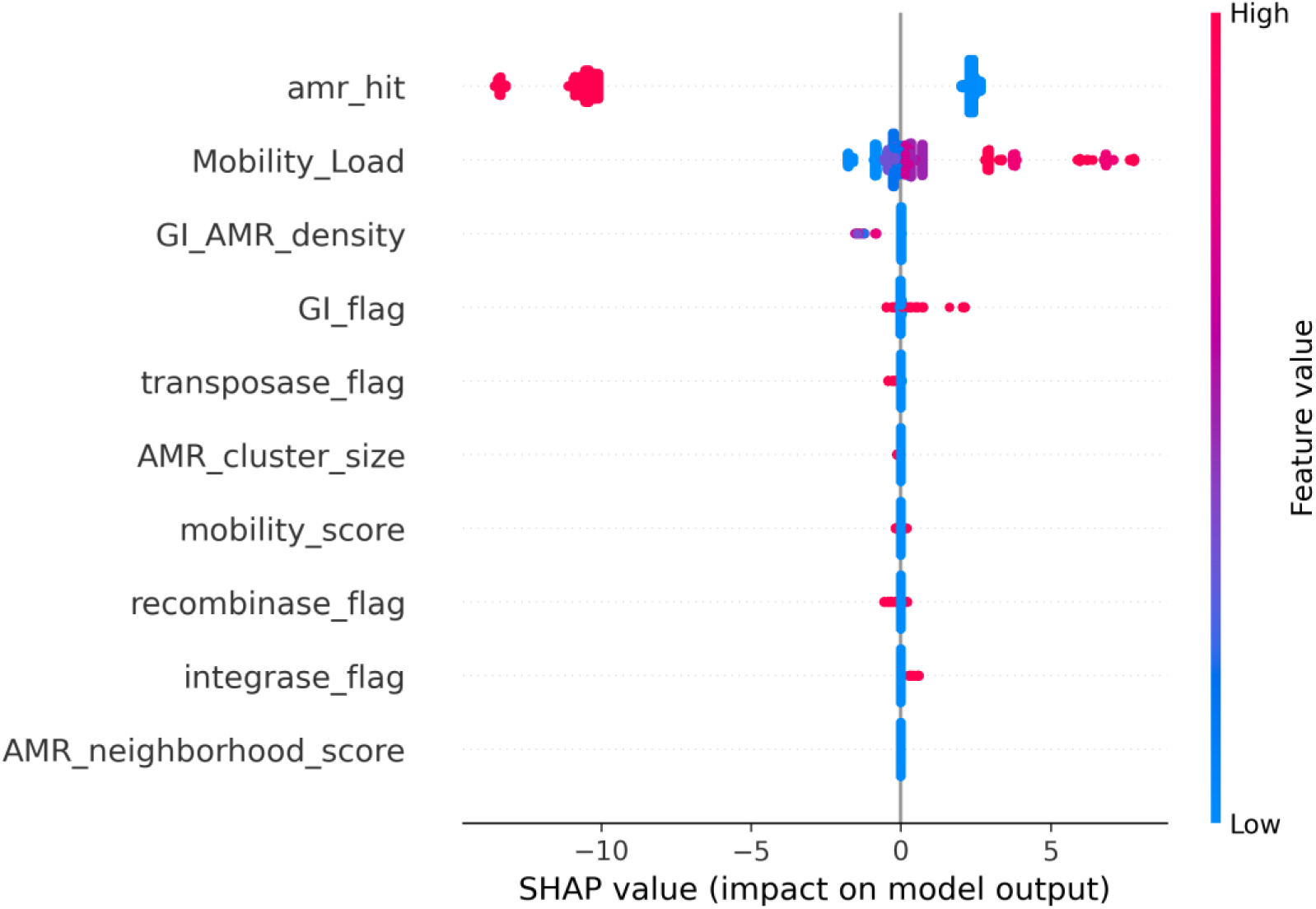
Model explainability. SHAP summary (beeswarm) plot showing the top cassette- and context-level features contributing to the model output under genome-grouped evaluation. Points represent per-sample SHAP values (impact on the prediction), with color indicating feature value (low to high). **Positive SHAP values increase the predicted probability of the Pathogenic class (as defined in the model), whereas negative SHAP values decrease it.**

### Comparative genomics contextualization

Comparative genomics plots further contextualize these findings. Pathogenic genomes show higher prevalence of several canonical AMR determinants (Supplementary Figure S1), and representative loci illustrate cassette-level AMR cargo architectures in a closed reference genome (Supplementary Figure S2). These patterns reinforce that while resistance genes contribute to the broader accessory genome landscape, non-AMR mobilome architecture and GI-anchored cassette organization provide additional, non-redundant signal captured by Cassette2Vec-EC.

## Discussion

The cassette architecture identified here is compatible with the hypothesis that module-scale horizontal acquisition and IS-mediated rearrangements contribute to pathogenic lineages, in which large, multi-gene genomic modules rather than individual genes may be mobilized between lineages. Enrichment of IS110-family transposases is consistent with increased insertion-sequence activity and associated genomic rearrangements (18, 34); representative members including isbth13 were identified through eggNOG-mapper functional annotation of GI-localized genes, although this locus fell outside the top 20 training-set-derived mobilome panel (Table S2). Insertion sequences can promote recombination, mobilization, and reorganization of neighboring loci, which could facilitate transfer or reassembly of multi-gene modules. The recurrent appearance of related cassette architectures across pathogenic lineages, as detected through IslandViewer and stabilized by PIRATE neighborhood conservation, supports the interpretation that horizontal transfer of module-scale genomic segments is a plausible contributor alongside incremental gene acquisition, although direct transfer events were not observed in this cross-sectional dataset.

Several high-impact cassettes encode a 5’-nucleotidase, an enzyme implicated in nucleotide metabolism and modulation of extracellular nucleotide pools. We hypothesize that extracellular ATP degradation could provide access to host-derived carbon and nitrogen sources while potentially dampening purinergic danger signaling by reducing extracellular ATP; these proposed metabolic and immunomodulatory roles are associative and require targeted functional validation in relevant infection contexts.

Enrichment of galactose metabolism pathways (KEGG map00052) (35) may reflect adaptation to carbohydrate and amino sugar availability in host-associated niches. We view this as a testable hypothesis that warrants targeted metabolomic and expression validation in vertebral and systemic infection contexts.

We interpret the predictive contribution of non-AMR loci in two non-exclusive ways: (i) some cargo genes may support host adaptation, persistence, or niche metabolism; and/or (ii) these loci act as high-fidelity structural ‘scaffolding’ markers that tag recurrent pathogenic genomic-island architectures. In either case, the signal reinforces that Cassette2Vec leverages module organization and GI context beyond resistance genes alone.

### Beyond Gene Presence: A Modular Genomics View

This work frames EC pathogenicity prediction as a structural-genomics problem: not only which genes are present, but how they are organized into cassette modules within predicted islands and mobility contexts. The key methodological contribution is enforcing genome-grouped evaluation while learning from cassette rows, which directly addresses a common failure mode in microbial ML studies (within-genome leakage).

Cassette2Vec-EC defines cassette units as local gene neighborhoods derived from IslandViewer-predicted genomic islands (21) and PIRATE-stabilized ortholog blocks (25), preserving physical adjacency, genomic-island context, and mobility signatures. Conceptually, this mirrors skip-gram or Word2Vec-style embedding logic (31, 32), where genes that co-occur within a cassette provide contextual information analogous to words within a sentence. By filtering ABRicate-identified AMR genes prior to embedding (22), the model is trained to learn mobility and architecture rather than resistance per se.

From a systems-microbiology perspective, cassette units provide a modular representation compatible with broader concepts such as pangenome graphs and modular evolution: genes that repeatedly co-occur within islands can be treated as transferable subgraphs whose presence, arrangement, and context influence host adaptation. Although demonstrated in EC, the Cassette2Vec architecture is portable to other bacteria (e.g., *Enterococcus faecium*, *Escherichia coli*, *Salmonella*) with minimal changes: (i) replace organism-specific annotation databases; (ii) tune cassette neighborhood definitions to genome architecture; and (iii) retrain under genome-grouped evaluation using the same GI/context feature schema.

Translationally, the framework supports practical near-term outputs: (i) ranked lists of high-risk genomes for targeted monitoring; (ii) cassette-level explanations to prioritize modules for wet-lab validation; and (iii) junction-based diagnostic designs anchored at recurrent cassette boundaries. These outputs align with industry surveillance needs and antibiotic-sparing control strategies.

## Limitations

This study has several limitations. First, the cohort is restricted to *Enterococcus cecorum* genomes sampled from poultry production systems, and generalization to other *Enterococcus* species, hosts, management systems, or geographic regions remains to be established. Second, available genomes may reflect sampling and ascertainment biases (e.g., overrepresentation of outbreak investigations, specific integrators, or production settings), and phenotype labels derived from metadata are inherently imperfect; isolates sampled from disease contexts may include bystanders, and “commensal” isolates may harbor pathogenic potential that is not expressed under the sampled conditions. Third, assembly-quality variation (e.g., contig fragmentation and N50 differences) can influence genomic-island prediction and cassette continuity: fragmented assemblies may split islands or interrupt gene neighborhoods at contig boundaries, potentially attenuating inferred structural and mobilome signals. Fourth, calibration is imperfect particularly at high predicted risk and varies across folds; thus, probability outputs should be interpreted as decision-support scores rather than definitive absolute risks unless post hoc recalibration and prospective validation are performed in the intended deployment setting.

Fifth, we did not perform external validation on a fully independent cohort outside the study corpus. However, key cassette and mobilome signatures identified here have been transferred to our industry partner (Cobb-Vantress) for prospective evaluation in ongoing surveillance programs; results from those deployments will be reported separately once sufficient follow-up data accrue. Finally, SHAP-identified cassettes and GI-associated features are associative and hypothesis-generating and should not be interpreted as causal determinants of pathogenicity without experimental validation. Long-read sequencing will enable validation of IslandViewer-predicted GI boundaries and direct confirmation of cassette contiguity inferred from short-read assemblies. A related concern is that performance may partially reflect phylogenetic clustering or BioProject-structured sampling (e.g., outbreak-associated clades) rather than cassette architecture alone. We therefore interpret portability claims conservatively until stratified evaluations (by BioProject/lineage/geography/year) and leave-one-source-out validation are completed. Because GI calling can vary across tools and assembly quality, cassette boundaries may shift with alternate GI predictors, GI-length thresholds, or contiguity heuristics; future work will quantify robustness to these alternate cassette definitions. Similarly, the ≥5 kb minimum island size threshold focuses on structurally meaningful mobile regions but excludes smaller horizontally acquired elements; sensitivity analyses using alternative thresholds (e.g., ≥3 kb and ≥10 kb) are planned to quantify effects on cassette inventory, feature distributions, and model performance. Together, these results indicate that the framework remains robust under fragmented assemblies so long as key GI anchors and neighborhood context are preserved, supporting practical deployment in surveillance settings where assembly quality varies.

## Future Directions

In addition, we will pursue true out-of-sample validation via leave-one-BioProject (or leave-one-lineage) out evaluation and external prospective cohorts when available, including applying the framework to newly sequenced isolates collected from new farms, time periods, and regions, enabling assessment of both discrimination and probability calibration under prospective surveillance conditions. These studies will also support stratified analyses by location, year, and lineage to quantify generalizability and identify settings where recalibration or retraining may be required.

Methodologically, we aim to move beyond vectorized representations toward graph-based learning, treating the accessory genome as a structured network in which cassette neighborhoods correspond to reusable subgraphs. Pangenome-graph-aware models could incorporate adjacency, ordering, and modular boundaries more directly, while preserving interpretability at the module level.

Translationally, we are developing junction-PCR assays targeting conserved boundaries between mobility-associated anchors (e.g., integrase/transposase/recombinase loci) and recurrent cargo modules, enabling rapid screening for high-risk architectures without requiring full genome assembly. Candidate junction targets will be prioritized from recurrent high-impact cassette archetypes and refined using long-read assemblies to validate cassette contiguity and boundary conservation prior to deployment in routine monitoring workflows.

## Conclusions

Taken together, these results support the idea that integrating structural genomics with machine learning is a practical next step beyond traditional pangenomic representations that emphasize gene presence-absence alone. In this paradigm, genomic architecture and genomic-island context provide informative biological signal that can be leveraged for prediction and hypothesis generation, alongside gene content.

Cassette2Vec-EC provides a cassette- and genomic-island-aware structural representation evaluated under genome-grouped validation, with strong predictive performance and interpretable cassette-level explanations. By linking genome-level risk estimates to GI-anchored modules, this framework supports actionable genomic surveillance of pathogenic *Enterococcus cecorum* and offers a transferable approach that can be adapted to other bacterial pathogens and production systems. Although demonstrated in *E. cecorum*, Cassette2Vec is organism-agnostic and can be applied to other bacteria in which genomic islands and modular horizontal transfer shape evolution, by re-running annotation/GI calling and retraining under the same genome-grouped protocol.

## Data Availability

All genome assemblies analyzed in this study are publicly available from NCBI under the following BioProjects: PRJDB18532, PRJEB32890, PRJEB33338, PRJEB50262, PRJEB50945, PRJEB54905, PRJEB57096, PRJNA1032545, and PRJNA1224097. Curated cohort metadata (genome identifiers, commensal/pathogenic labels, and assembly/annotation-derived summary metrics) are provided as Supplementary Data S1

(Supplementary_Data_S1_Cohort_Metadata.xlsx). Mobilome signature tables and stability frequencies are provided as Supplementary Data S2

(Supplementary_Data_S2_Mobilome_Signature_20gene.xlsx). Per-genome genomic-island (GI) burden summaries are provided as Supplementary Data S3

(Supplementary_Data_S3_per_genome_GI_burden.xlsx). Where available, raw sequencing reads are accessible via the SRA accessions linked within each BioProject record.

## Code Availability

Cassette2Vec-EC source code is available at GitHub (https://github.com/RushiLagad/Cassette2Vec; tag v1.0.0; commit 278a146; accessed 2026-02-02). A versioned archive corresponding to this release is deposited on Zenodo (v1.0.0 DOI: https://doi.org/10.5281/zenodo.18529389). The concept DOI, which resolves permanently to the latest release for all future versions, is https://doi.org/10.5281/zenodo.18529388; readers are encouraged to cite the version DOI for full reproducibility. The repository includes scripts for preprocessing/annotation, genomic-island parsing, PIRATE-based ortholog processing, cassette construction and feature engineering, model training/evaluation under genome-grouped cross-validation, and SHAP analyses used for interpretability. All analyses are reproducible using the provided run scripts and configuration files; the repository includes environment specifications and fixed random seeds, with step-by-step reproduction instructions in the README.

## Acknowledgments

This research was primarily supported by the Arkansas Research Alliance (ARA) Impact Grant Program, with additional support from Cobb-Vantress Genetics LLC. We thank the Department of Animal Science and the Center for Agricultural Data Analytics (CADA), University of Arkansas, for institutional support and guidance. This research used resources of the Arkansas High Performance Computing Center, which is funded through multiple National Science Foundation grants and the Arkansas Economic Development Commission.

## Author Contributions

Rushikesh R. Lagad designed and implemented the Cassette2Vec-EC pipeline, performed all genomic analyses and machine learning experiments, generated the visualizations, and drafted the manuscript. Shakil Rafi contributed to machine learning methodology, model validation, and data science implementation, and reviewed the manuscript critically. Aranyak Goswami conceived and directed the study, acquired funding, provided scientific oversight throughout, and contributed to manuscript writing and final drafting. All authors read and approved the final manuscript.

## Abbreviations

AMR: Antimicrobial resistance
AUPRC: Area under the precision-recall curve
AUROC: Area under the receiver operating characteristic curve
CARD: Comprehensive Antibiotic Resistance Database
COG: Cluster of Orthologous Groups
DIMOB: Dinucleotide bias and mobility (IslandPath-DIMOB module)
FASTA: Text-based format for nucleotide or protein sequences
FAA: Amino acid FASTA file
FNA: Nucleotide FASTA file
GFF3: General Feature Format version 3
GI: Genomic island
HMM: Hidden Markov Model
IS: Insertion sequence
KEGG: Kyoto Encyclopedia of Genes and Genomes
MGE/MGEs: Mobile genetic element(s)
MLS: Macrolide-lincosamide-streptogramin resistance
N50: Assembly contiguity statistic (length at which 50% of the genome is contained in contigs ≥ N50)
NCBI: National Center for Biotechnology Information
PCA: Principal component analysis
PIRATE: Pangenome Integration of Reproducible Annotation Tools for Evolution
SHAP: SHapley Additive exPlanations
SIGI: SIGI-HMM composition-based genomic island predictor
UMAP: Uniform manifold approximation and projection
VF: Virulence factor
WGS: Whole-genome sequencing
XGBoost: Extreme Gradient Boosting (tree-based ML algorithm)

## Supplementary Figures

**Figure S1.**
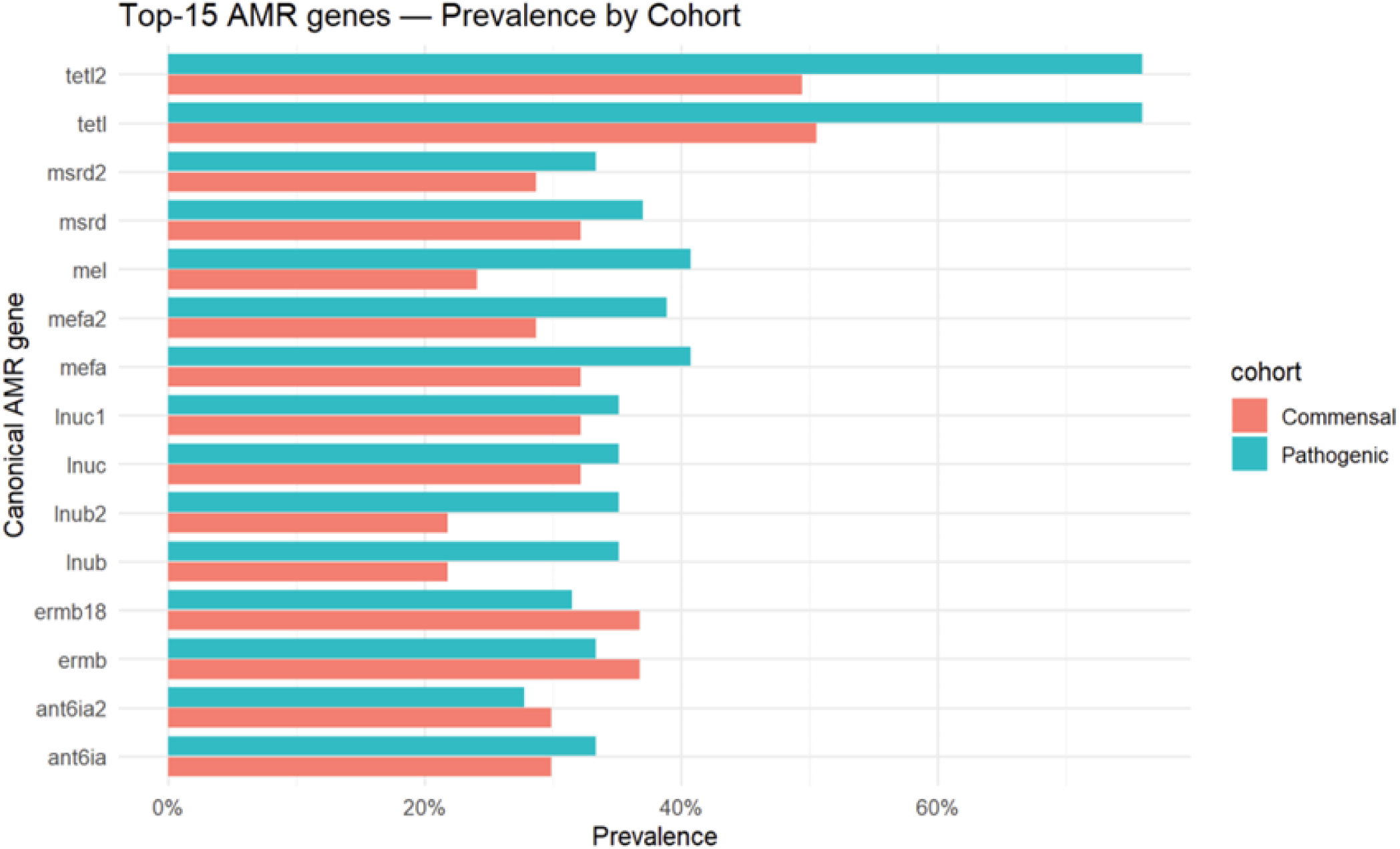
Top-15 AMR genes prevalence by cohort (commensal vs pathogenic).

**Figure S2.**
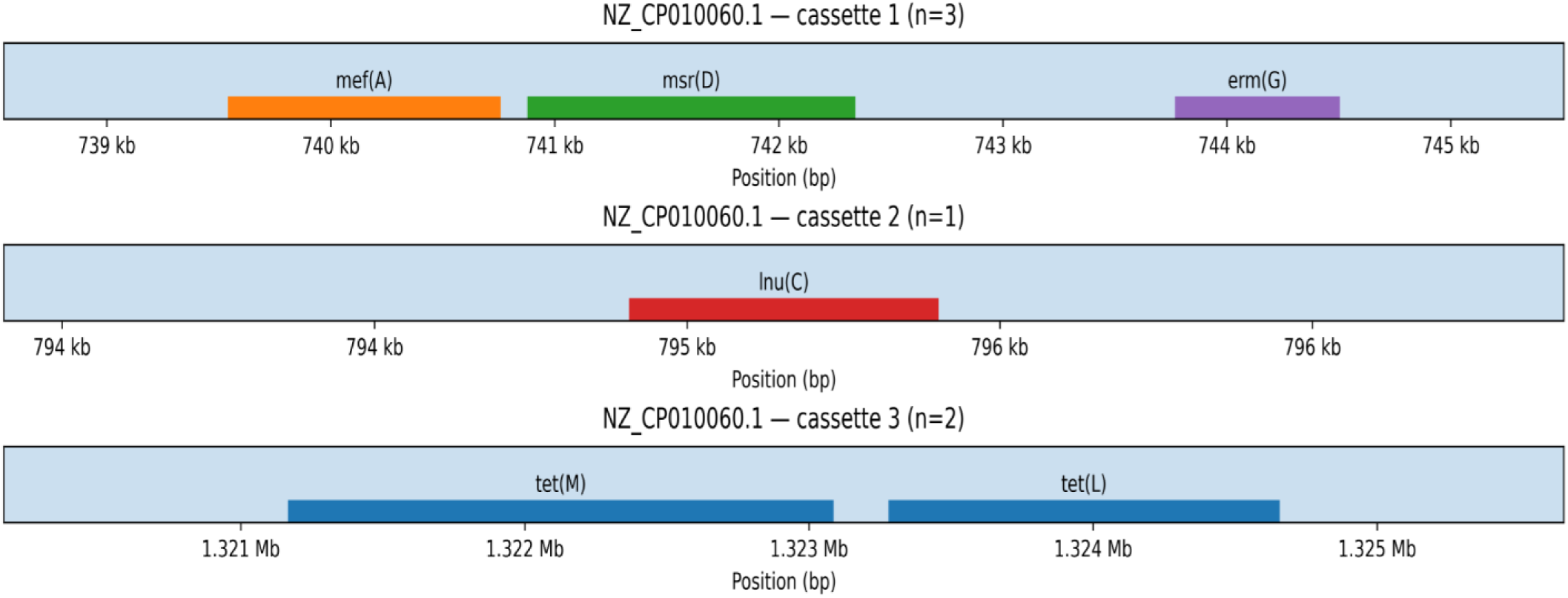
Representative AMR cassette architectures in reference genome NZ_CP010060.1 (example loci).

---

**Table S1.** Full cohort metadata (n = 145 genomes) provided as Supplementary Data S1 (Supplementary_Data_S1_Cohort_Metadata.xlsx). A brief excerpt is shown below for formatting reference.

| Genome ID | Class | Host/source | Country/Region | Year | Assembly accession | Assembly quality (N50/contigs) |
| --- | --- | --- | --- | --- | --- | --- |
| GCF_000379745.1 | commensal | poultry | NA | NA | GCF_000379745.1 | NA / 45 |
| GCF_000407565.1 | commensal | poultry | NA | NA | GCF_000407565.1 | NA / 5 |
| GCF_000492155.1 | commensal | poultry | NA | NA | GCF_000492155.1 | NA / 2 |
| GCF_001020435.1 | commensal | poultry | NA | NA | GCF_001020435.1 | NA / 23 |
| GCF_001020445.1 | commensal | poultry | NA | NA | GCF_001020445.1 | NA / 24 |
| GCF_001021675.1 | commensal | poultry | NA | NA | GCF_001021675.1 | NA / 24 |
| GCF_001021745.1 | commensal | poultry | NA | NA | GCF_001021745.1 | NA / 40 |
| GCF_001021755.1 | commensal | poultry | NA | NA | GCF_001021755.1 | NA / 50 |
| GCF_001021795.1 | commensal | poultry | NA | NA | GCF_001021795.1 | NA / 40 |
| GCF_001021815.1 | commensal | poultry | NA | NA | GCF_001021815.1 | NA / 53 |

**Table S2.**
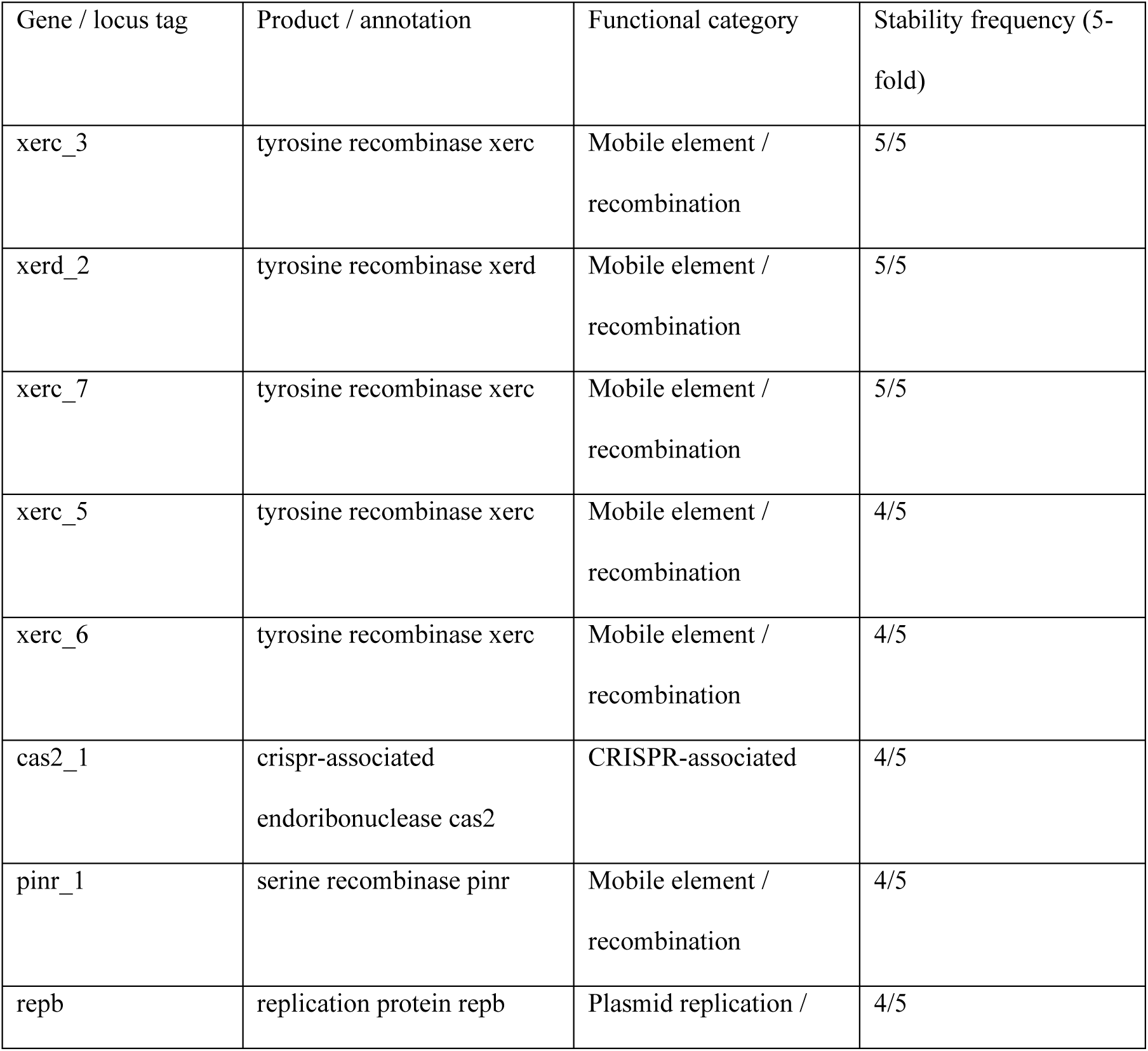
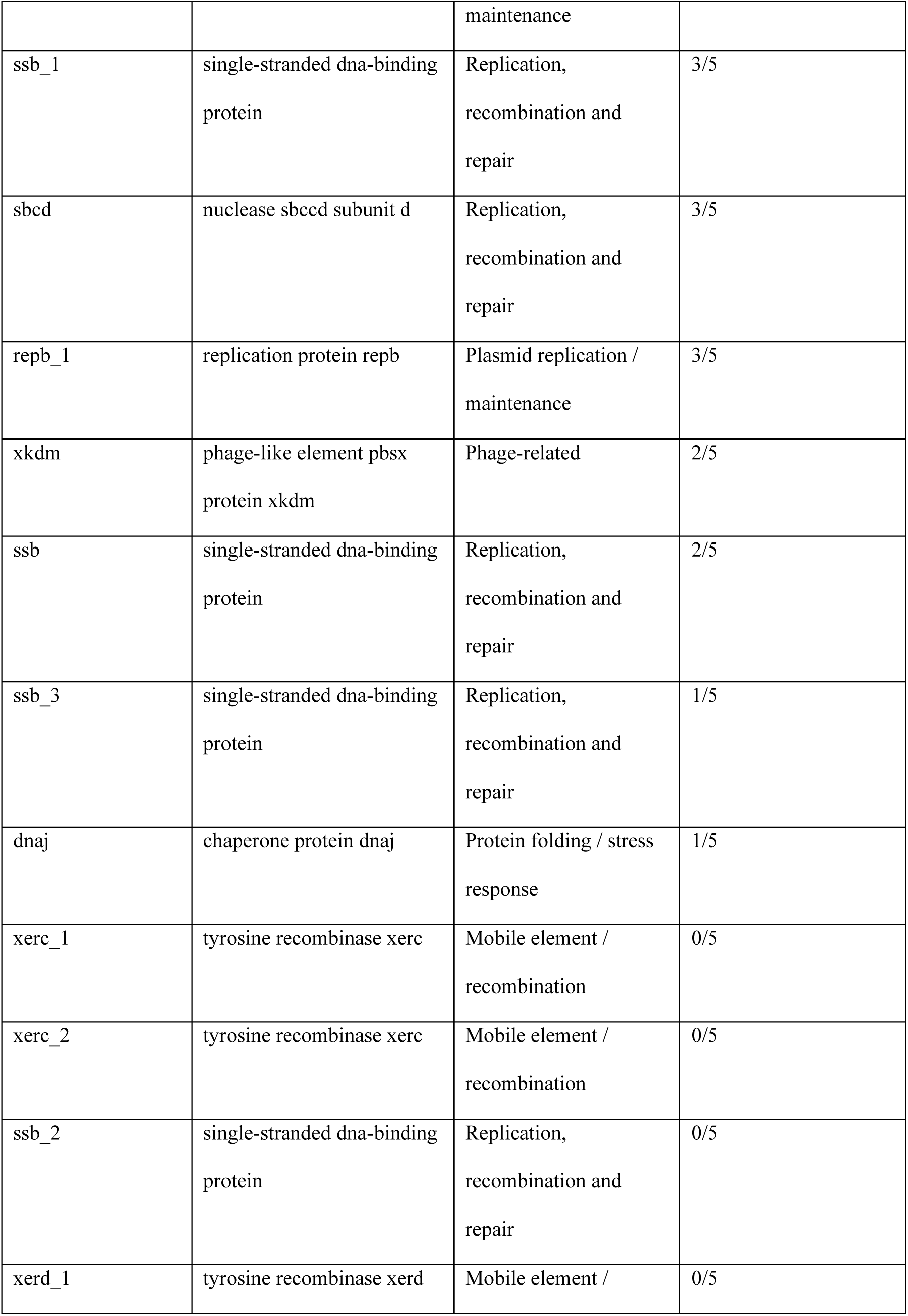
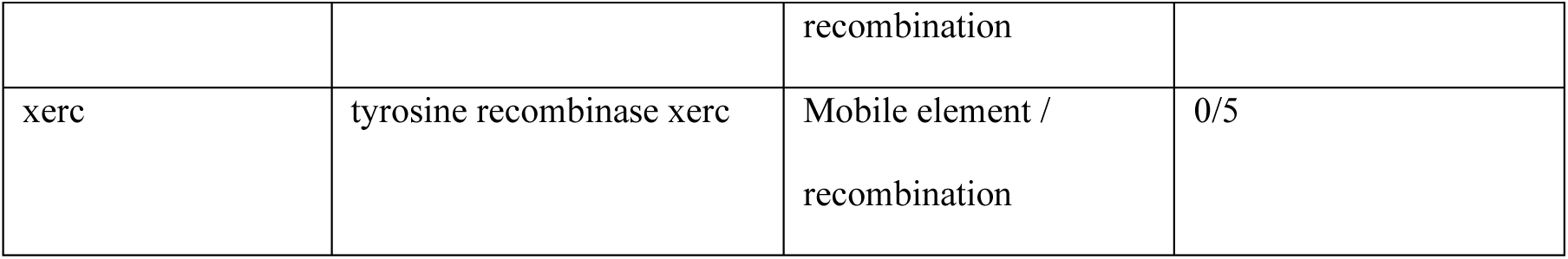
20-gene non-AMR mobilome signature with functional annotations and stability across outer-fold training splits. Stability frequency reports, out of 5 outer training splits, how often each gene ranked among the top 10 pathogenic-enriched signature loci based on log2 odds ratio of GI-localized presence.

**Table S3.** Per-genome genomic-island (GI) burden summaries (IslandViewer-available subset) provided as Supplementary Data S3 (Supplementary_Data_S3_per_genome_GI_burden.xlsx). A brief excerpt is shown below for formatting reference.

| Genome ID | Class | Predicted GI count | Total GI length (kb) |
| --- | --- | --- | --- |
| GCF_000379745.1 | commensal | 15 | 267.36 |
| GCF_000407565.1 | commensal | 16 | 241.57 |
| GCF_000492155.1 | commensal | 15 | 238.45 |
| GCF_001020435.1 | commensal | 11 | 204.29 |
| GCF_001020445.1 | commensal | 10 | 265.86 |
| GCF_001021675.1 | commensal | 12 | 252.84 |
| GCF_001021745.1 | commensal | 11 | 214.66 |
| GCF_001021755.1 | commensal | 11 | 187.06 |
| GCF_001021795.1 | commensal | 14 | 237.13 |
| GCF_001021815.1 | commensal | 12 | 299.81 |

**Table S4.**
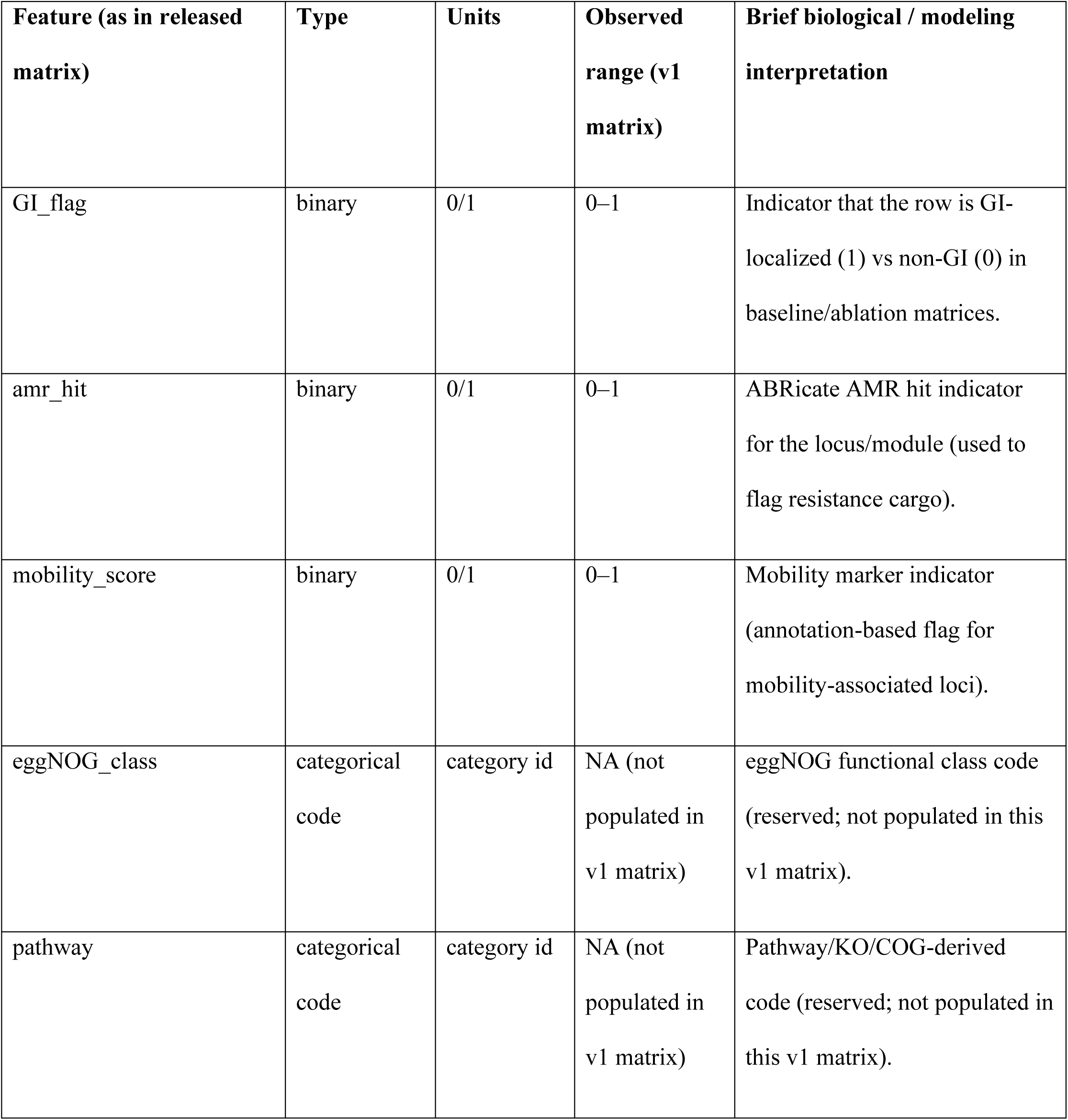
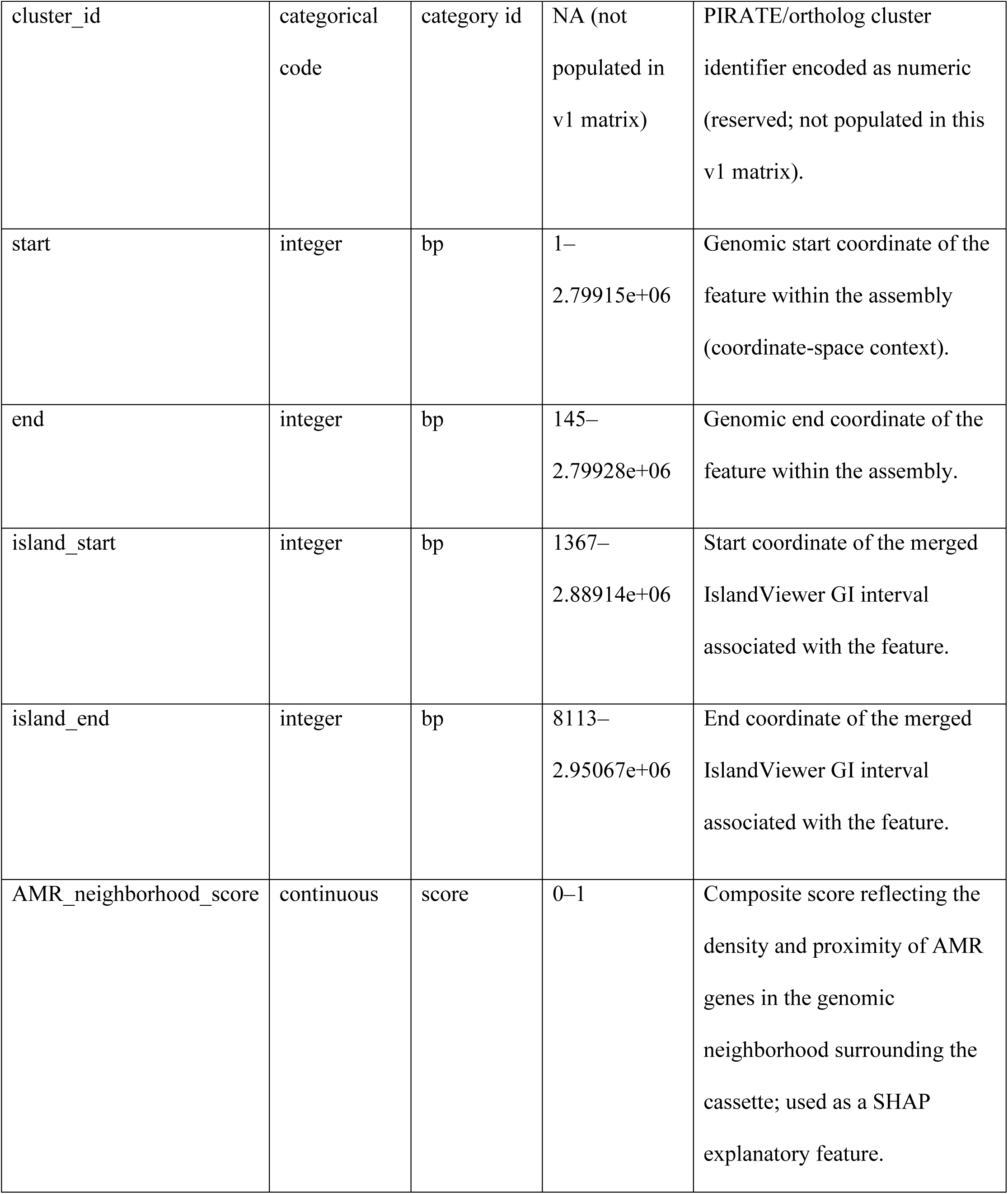
Cassette2Vec-EC “Full (all numeric)” 20-feature input schema. Cassette2Vec-EC “Full (all numeric)” model input feature definitions (n = 20). This table lists the complete set of cassette-level numeric inputs used by the Full model, with feature names exactly as provided in the released matrix, their data type, units, observed ranges in the analyzed cohort, and a brief biological/modeling interpretation. Rows 1-10 are shown here as a formatting reference; all 20 feature definitions, including the three reserved schema fields (eggNOG_class, pathway, cluster_id) marked is_populated = FALSE, are provided in Supplementary Data S4 (Supplementary_Data_S4_Cassette2Vec_Full20_Feature_Definitions.xlsx).

